# Different tastes for different individuals

**DOI:** 10.1101/009357

**Authors:** Kohei Fujikura

## Abstract

Individual taste differences were first reported in the first half of the 20th century, but the primary reasons for these differences have remained uncertain. Much of the taste variation among different mammalian species can be explained by pseudogenization of taste receptors. In this study, by analyzing 14 ethnically diverse populations, we investigated whether the most recent disruptions of taste receptor genes segregate with their intact forms. Our results revealed an unprecedented prevalence of segregating loss-of-function (LoF) taste receptor variants, identifying one of the most pronounced cases of functional population diversity in the human genome. LoF variant frequency (2.10%) was considerably higher than the overall mutation rate (0.16%), and many humans harbored varying numbers of critical mutations. In particular, molecular evolutionary rates of sour (14.7%) and bitter receptors (1.8%) were far higher in humans than those of sweet (0.02%), salty (0.05%), and umami (0.17%) receptors compared with other carnivorous mammals although not all of the taste receptors genes were identified. Many LoF variants are population-specific, some of which arose even after the population differentiation, but not before divergence of the modern and archaic (Neanderthal and Denisovan) human. Based on these findings, we conclude that modern humans might have been losing their taste receptor genes because of high-frequency LoF taste receptor variants. Finally I actually demonstrated the genetic testing of taste receptors from personal exome sequence.

## Introduction

Taste is a primitive sense that helps recognize and distinguish key dietary components (1–3). Taste perception affect nutrient intake and appetite, and thus taste decrements can lead to food poisoning and over-exposure to hazardous chemicals (1,4). The sensation of taste is classified into five prototypical categories: sweet, bitter, sour, salty, and umami (1–3,5–7). In addition recent compelling evidence raises the possibility of an additional sixth and seventh taste modality devoted to the perception of calcium (8,9) and lipids (10,11). We all perceive these tastants through sensory organs called taste buds (2,3,5-7). Humans have approximately 10,000 taste buds, primarily located on the tongue with a few on the palate and throat (1).

The anatomical units of taste detection are taste receptor cells (TRCs) (1–3). TRCs are distributed among different lingual papillae and palate epithelia, and are involved in recognizing various taste stimuli (1–3). The sensation of taste is produced when stimuli in the mouth react with specific taste receptors (1–3,5–7). Taste coding at the peripheral cells may rely on straightforward labeled lines (that is, sweet, bitter, sour, salty, and umami signals) to transform tastant quality into neural signals through the glossopharyngeal and facial nerve (1,3). Taste in conjunction with olfaction is perceived as flavors or other substances (1–3).

During the last 10 years, the discovery and characterization of mammalian taste receptors has progressed tremendously (1–3,6,7). The recent identification of cells and receptors that mediate the prototypical tastes has generated powerful molecular tools that can be used to devise tests to establish the basis of taste coding at the periphery (12–18); tastant-induced activity were measured using nerves innervating the buds in various gene-knockout (KO) mice (2,15) or receptor-expressing cells (16–18), and diphtheria toxin A fragment (DTA)-mediated cell ablation studies were performed to uncover the specific taste stimuli *in vivo* (19).

“Why do some people have different taste recognition thresholds compared with others?” “Do they have different receptor densities within their body?” These have been controversial issues since the early 20th century and the reasons for individual taste differences remain uncertain (20–26). Recent studies have raised the possibility that most genotype–phenotype association studies are based on rare gene disruptions arising during mammalian evolution (27,28). In fact, some taste receptors also seem to be affected by mutations during mammalian evolution (6,7,12,13,29-31). For example, the *hT2R43* W35 allele makes people most sensitive to the bitterness of the natural plant toxins, aloin and aristolochic acid, and an artificial sweetener, saccharin, compared with S35 allele (31). In addition, a closely related gene’s *hT2R44* W35 allele is also sensitive to saccharin (31). People who do not possess these allele do not taste the bitterness of these compounds at low micromolar concentrations (31). Another well-known example is *hTAS2R38*. African populations have higher levels of genetic, geographic and phenotypic diversity at the *TAS2R38* locus (32). *TAS2R38* P49A, A262V and V296I have clear effects on the phenylthiocarbamide (PTC) sensitivity in humans (33–35). These polymorphisms illustrate the influence of recent genetic variation on a common trait. However none of the genetic variants explain the whole spectrum of interindividual differences of taste sensitivity.

We postulated that individual taste differences were predominantly caused by loss-of-function (LoF) variants of taste receptor genes. In this study, we investigated whether the most recent of these disruptions may still segregate with the intact forms by analyzing 14 ethnically diverse populations. The recently sequenced completed genomes for many modern humans provided the opportunity for reconstruction of the currently known LoF variants (37,38). Our results revealed an unprecedented prevalence of segregating LoF variants, one of the most pronounced cases of functional population diversity in the human genome.

## Results

### Profiles of human taste receptor gene variations reveal striking individuality

The representative taste receptors are composed of more than 50 coding regions distributed in clusters over most chromosomes in mammals (5). In mice, taste receptor pseudogenes comprise 15% of this gene range; in humans, a fraction roughly two times larger appears to be inactivated (5). Of more than 50 taste receptor genes, more than half appear to have been pseudogenized by mutations during mammalian evolution (5). Extreme diminution of the functional taste receptor repertoire was a relatively recent genomic process and is probably still ongoing (5). Therefore, we conjectured that a substantial fraction of modern human taste receptors may segregate between intact and pseudogene forms. Isolated cases of segregating mutations in taste receptors have been reported (6,7,12,13,29–31).

We focused on three kinds of LoF variants expected to correlate with complete loss of function of the affected transcripts: one stop codon–introducing (nonsense) or two (5’ and 3’) splice site disrupting single-nucleotide substitutions. We searched 1,000 genomes (37) and NHLBI databases (38) for variations with the potential to affect the protein integrity of taste receptor genes (fig. 1 and table S1). Target receptor genes are as follows: *TAS1R1*, *TAS1R2*, *TAS1R3*, *TAS2R1*, *TAS2R3*, *TAS2R4*, *TAS2R5*, *TAS2R7*, *TAS2R8*, *TAS2R9*, *TAS2R10*, *TAS2R13*, *TAS2R14*, *TAS2R16*, *TAS2R19*, *TAS2R20*, *TAS2R30*, *TAS2R31*, *TAS2R38*, *TAS2R39*, *TAS2R40*, *TAS2R41*, *TAS2R42*, *TAS2R43*, *TAS2R45*, *TAS2R46*, *TAS2R50*, *TAS2R60*, *PKD1L3*, *PKD2L1*, *CD36*, *ENaCa*, *ENaCd*, *HCN1*, *HCN4*, *TRPM5*, and *CALHM1* (table S1). I did not analyze the putative taste receptor genes and perform structural similarity searches. Taste receptor are different from olfactory receptors (ORs) in that taste receptors are not composed of single gene family (1–3). Our data matrix included approximately 8,000 individuals from six ethnic origins (table S2). In this largest realm, we extrapolated that the number of major segregating taste receptor LoF variants in the entire human genome had to be at least 24 of which 18 were expected to have a major allele frequency of >0.05% (Figure 2A). The average LoF variant frequency among known taste receptor genes (2.10%; TAS2R (18) 24, = 1.76%; PKD-like (PKD1L3+PKD2L1 (2,3) = 14.7%) was extremely higher than the overall mutation rate in the human genome (approximately 0.16% (calculated from LoF variants caused by nucleotide substitution; this proportion also includes both LoF frequency of taste and smell receptor genes) (38)) (fig. 2B). Except for complete pseudogenes, mutation rates of taste receptor genes exceeded those of olfactory receptor (OR) genes (approximately 1.41%; recalculated from 367 functional ORs because more than half genes analyzed in a past study (39) have already been proved to be complete pseudogenes; If we calculate the mutation frequency of pseudogenes, their MAF is regarded as 100% and mutation frequency become quite high).

**Figure 1.**
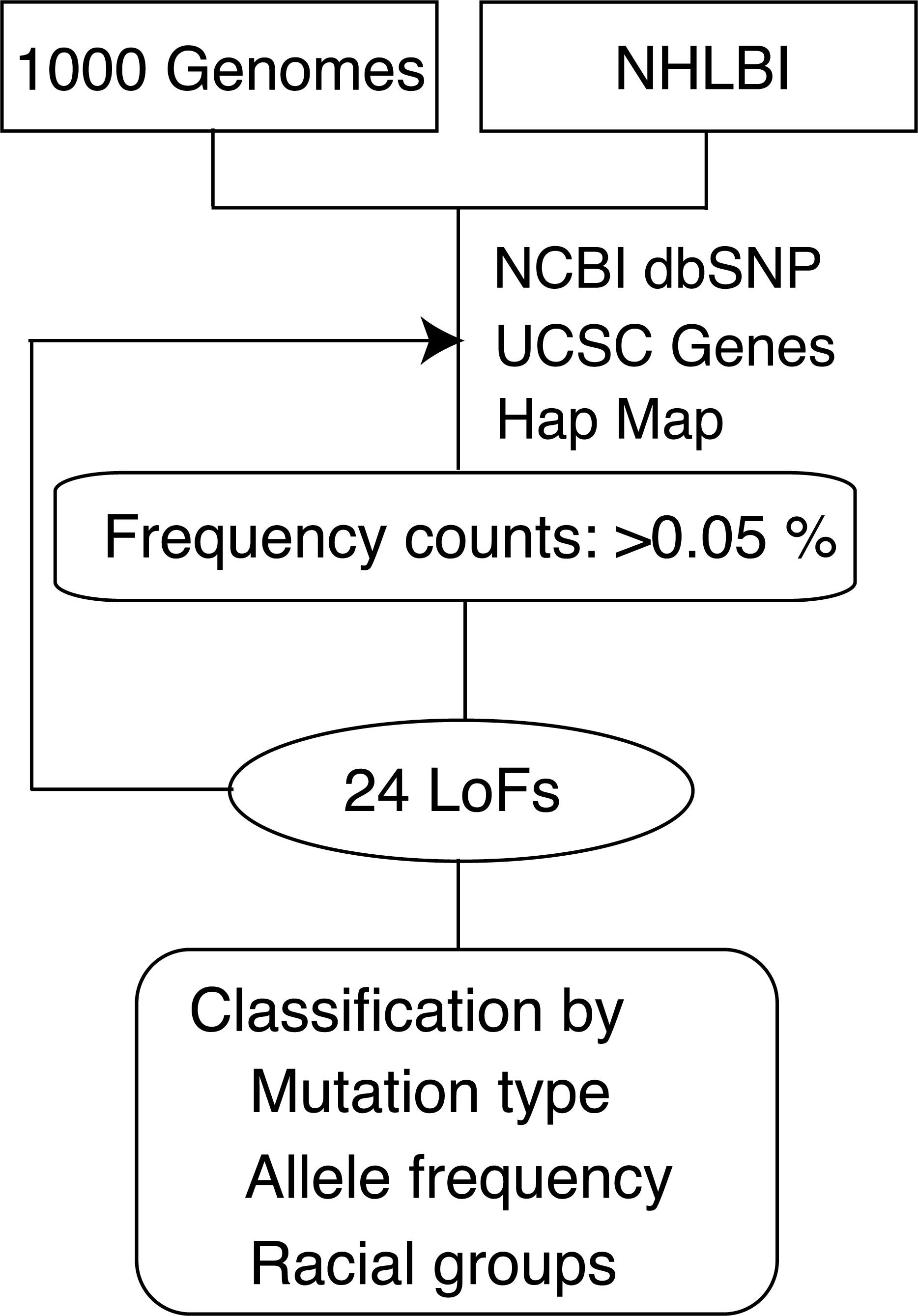
Strategy for identification of deleterious mutations within taste receptor genes. A flow chart used to identify the sequence variations that have harmful influence on human taste senses shows 25 LoF variants within taste receptor genes. Several platforms (NCBI dbSNP, UCSC Genes, and Ensemble variation) were used to access the validity of variants and examine previous gene annotations.

**Figure 2.**
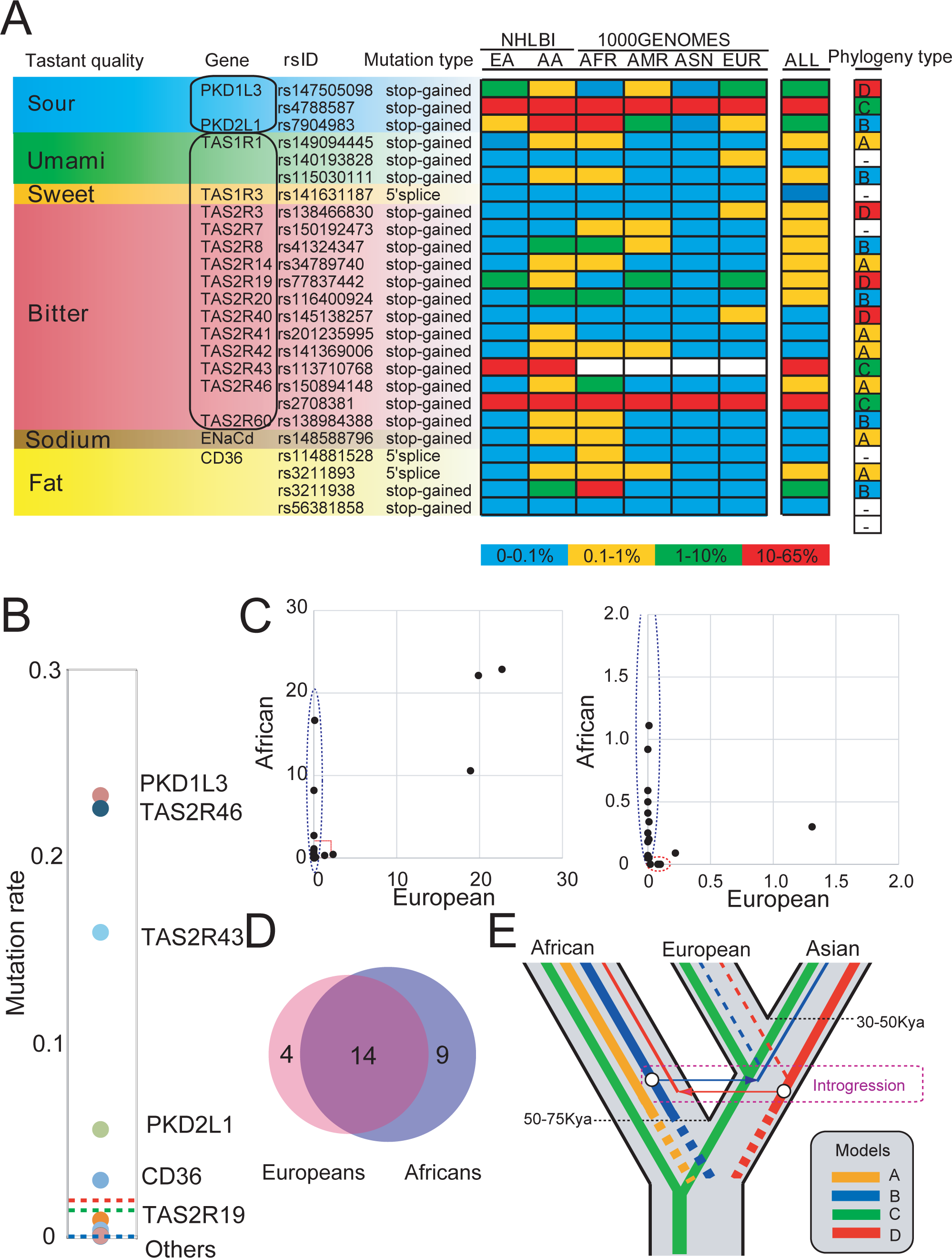
Loss-of-function variants of taste receptors in major ethnic groups. (A) Allele frequency of LoF variants of 18 taste genes in European (EA) and African (AA) American (NHLBI), and African (AFR), Ad mixed American (AMR), East Asian (ASN) and European (EUR) (1000 Genomes). Colored boxes indicate the frequency of LoF variants (blue, 0-0.1 %; yellow, 0.1-1 %; green, 1-10 %; red, 10-65 %).Taste genes are ordered according to their tastant quality and functions. dbSNP ID and disruption type is described. (B) Distribution of LoF frequency of each tastant genes. Blue, Green and red lines indicate average LoF frequency of taste receptors (2.10 %), olfactory receptors (1.41 %) and overall genes of human genome (∼0.16 %), respectively. (C) Comparison of MAF (%) distribution of taste receptor genes between African and European. Vertical and horizontal axis indicate MAF in African and European, respectively. Right panel is enlarged view of the red box at the left panel. Blue and red dotted circle indicate the population-specific or enriched variants. (D) Venn diagram showing the number of shared and unique LoF variants for taste receptor genes between European American and African American. (E) Hypothetical origins of different taste between ethnic groups. Divergence of taster and nontaster alleles before and after race divergence is indicated as four lines (phylogeny type A, B, C, D).

In theory, a combination of these genes could give rise to an enormous series of individual differences in taste perception; each examined individual had a unique genotypic pattern. The average number of LoF sites in TAS1R (2,3), TAS2R (18) and PKD-like (PKD1L3+PKD2L1 (2,3)) receptors per individual was 0.00371, 0.422 and 0.294, respectively. Coupled with TAS and PKD-like genes, *CD36* (fat receptor (10,11)), and *ENaCd* (*SCNN1D*) (sodium channel (40–43))) were frequently lost in some populations (fig. 2A, table S2 and S3). In contrast, we confirmed that the other taste buds-specific or -enriched genes previously reported (44,45) had been rarely lost (forty five genes analyzed are described in supplementary note).

No novel taste receptor genes were acquired after the divergence of chimpanzees and humans. Using 1000 genome sequences, we confirmed that the three pseudogenized bitter receptors *TAS2R2P*, *TAS2R62P*, and *TAS2R64P* (46) have never been functionally restored by gain-of-function mutations in any human population. Our survey of LoF variants in taste receptor genes pointed out the possibility that modern humans might have continued to genetically lose the repertoire of receptor genes after species differentiation.

### Population-specific LoF variants in taste receptor genes

Because of physical barriers to migration, ethnic populations rarely interbreed as convenient in theoretical random models. Consequently populations from different ethnic groups often have different genetic backgrounds and therefore different frequencies of genetic polymorphisms (47). To examine the genetic distance of taste receptor genes among populations, we first investigated whether frequencies and distributions of mutations had regional and racial biases by constructing a map for relationships between mutation rates and gene locations (fig. S1 and table S2 and S3). Comparative analyses showed that patterns of nucleotide substitution rates varied substantially among different regions of the genome and among different ethnic populations (fig. S1 and table S2 and S3).

These results suggested that some taste receptor variants had different origins and emphasized the necessity to narrow the range of target sequences that are to be searched on the basis of the ethnic background. Of note, African samples provided a huge resource for discovering variant sites, whereas non-African individuals had significantly fewer variations (fig. 2C and D and table S2). Furthermore, the non-African diversity was largely a subset of the African diversity (fig. 2C and D and table S2).

By comparing alleles among individuals from various ethnic backgrounds, we further estimated the extent of differentiation in taste receptor genes before and after the divergence from African origins (48) approximately 50,000–75,000 years ago. We used a pair-wise proportions test (49), which is used for testing a null hypothesis stating that proportions in two populations are the identical. This is referred to as a z-test because the statistic is

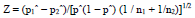

where p^ = (p_1_ + p_2_)/(n_1_ + n_2_), and the indices (1, 2) refer to the first and second line of the table.

We divided the LoF events of taste receptors into four phylogenies (types A, B, C, and D) based on a significant difference (*P* < 0.05) between populations, suggesting that humans might have always been losing their taste receptors, even after population differentiation (fig. 2A and E). Moreover, our results revealed that gene mutation flow from Africans to non-Africans, and vice versa, had occurred (fig. 2E). This evidence suggested that the general theory (50–52) that different evolutionary pressures, such as diets, toxins, and climates (energy consumption) have shaped the different chemosensory repertoires in mammals might be applicable to modern human populations.

We next used multivariate analysis to investigate country-by-country differences in taste LoF variants. To separate confused data sets to make several distinct classes, Hierarchical Ward’s analysis was used for the matrices of spectra from 14 ethnic groups to compare their profiles of taste receptor gene variations (fig. 3A, left panel; see also fig. S2). Hierarchical Ward’s method showed that 14 ethnic populations could be categorized into three general groups: African, Asian, and European–Hispanic. Hierarchical median algorithms (fig. S3) and non-Hierarchical clustering algorithms (k-means algorithms; fig. 3A, right panel) also supported these categories.

**Figure 3.**
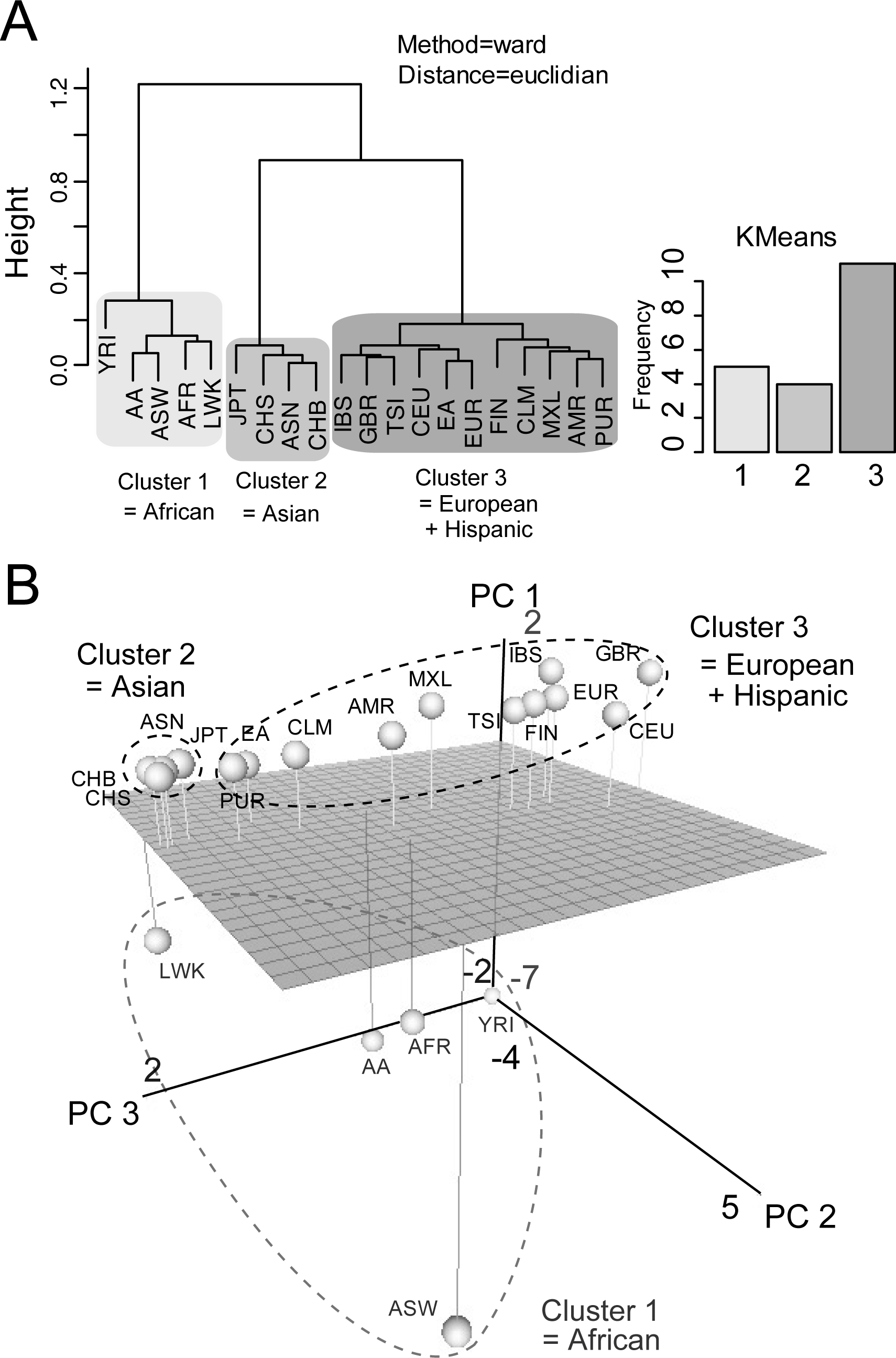
Multivariate analysis of genetic variants across 14+6 ethnic populations. (A) Hierarchical (dendrogram; ward’s algorithm) and Non-Hierarchical (bar graph; k-means algorithm) clustering produce three genetic clusters across 14+6 ethnic populations (ASW, American’s of African Ancestry in SW; CEU, Utah Residents (CEPH) with Northern and Western European ancestry; CHB, Han Chinese in Beijing; CHS, Southern Han Chinese; CLM, Colombian from Medellian; FIN, Finnish in Finland; GBR, British in England; IBS, Iberian population in Spain; JPT, Japanese in Tokyo; LWK, Luhya in Webuye; MXL, Mexican ancestry from Los Angeles; PUR, Puerto Rico from Puerto Rica; TSI, Toscani in Italia; YRI, Yorba in Ibadan) (AFR, African; ASN, East Asian; AMR, Ad mixed American; EUR, European; EA, European American; AA, African American). (B) PCA projection of samples taken from a set of 14+6 ethnic groups. Grey-scaled spheres correspond to Asian, African, European and Hispanic.

Furthermore, we used principal component analysis (PCA) to compare the LoF variants of taste receptor genes among various countries (fig. 3B). The three principal components (PC 1, 2 and 3) reflected the difference of taste LoF variants in these populations. These structure patterns were consistent between the two approaches (fig. 3A and B). However, when using the country-by-country approach, there were large amounts of differences in taste LoF variants among African, European and Hispanic populations (fig. 3B). In contrast, taste genetic structures differed only slightly among Asian populations (fig. 3B). The taste LoF patterns in Hispanics was genetically closer to those in Europeans than the other populations, and some Hispanic (PUR and CLM) and African (LWK) groups are similar to Asian populations (fig. 3B). East Asian groups, including Japanese and Chinese, had only a few LoF variants of bitter receptor genes (table S3). These results illustrated country-by-country genomic signatures for the taste LoF variants in humans and sufficient proof that taste receptor genes had evolved further in individual colonized areas.

### Independent origins of mutant alleles

Previous studies raise the possibility that different ethnic groups often shared same taste sensitivity (for example, sweet, salt, bitter and so on) to various compounds mediated by taste receptors (53–56). However, phylogenetic relationships among LoF taste receptor variants were, for the most part, consistent with the hypothesis that some LoF alleles had independently arisen at least twice between the two ethnic groups over the course of human evolution (fig. 2A). For example, both rs150894148 and rs2708381 caused stop-gain mutations in *TAS2R46* at almost the same position, although their origins were speculated to be different (rs150894148: phylogeny type A; rs2708381: phylogeny type C; fig. 2A).

The molecular evolutionary rates of sour and bitter receptor genes seems much higher than those of the other receptor genes in any ethnic group (fig. 4). In general sour or bitter tastes can be unpleasant, but–because many toxic substances taste sour or bitter―they can also be life-saving. My result raised the possibility that modern humans lacking sour and/or bitter taste receptors would not seemingly be at a significant disadvantage. At present not all of the taste related genes were analyzed, and thus confirmation will have to wait until further study is conducted. Characterizing these patterns could facilitate us to unveil the evolutionary pressure acting on modern humans.

**Figure 4.**
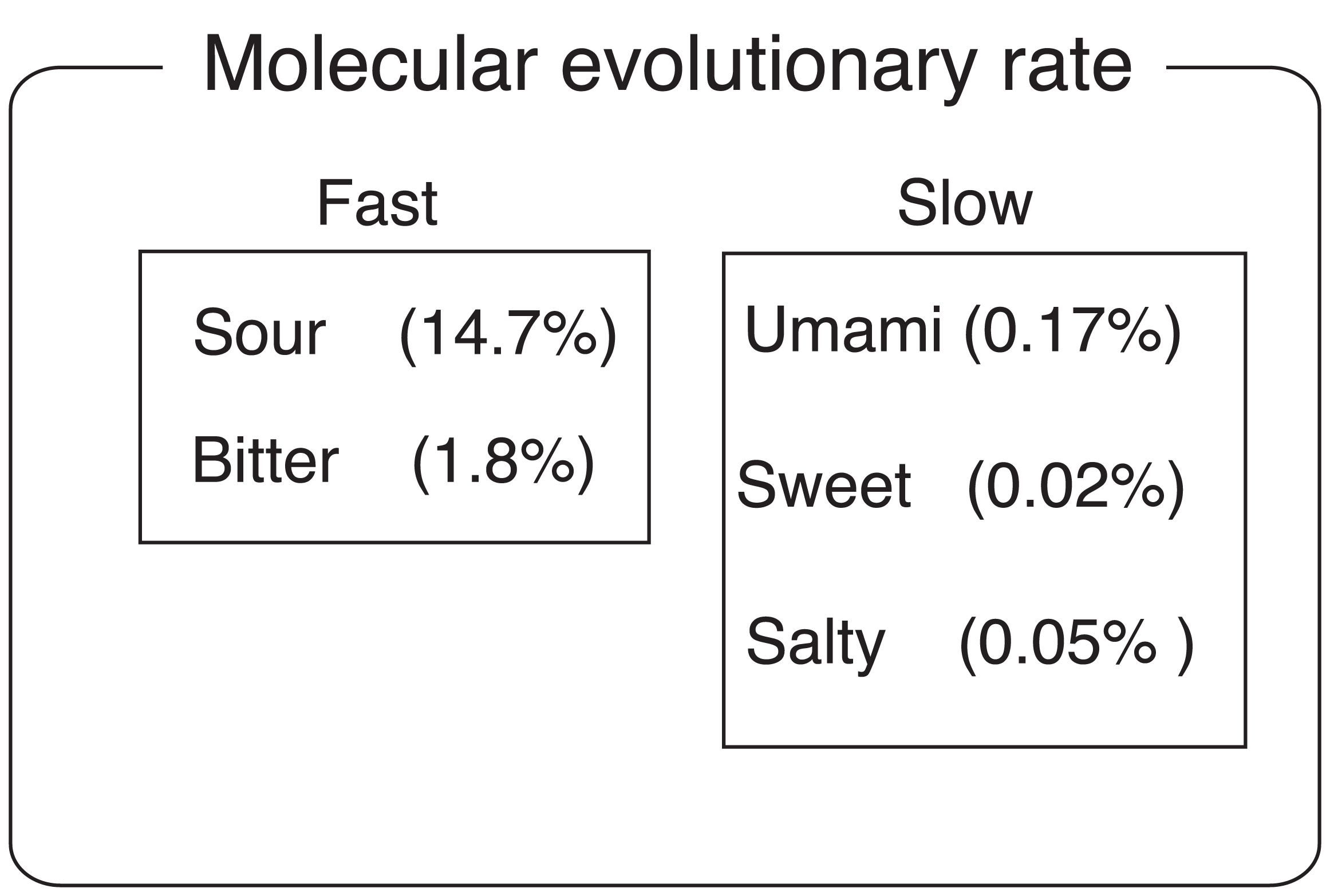
Molecular evolutionary rate of five taste senses in modern humans. Average LoF mutation frequency of taste receptor genes of five senses (sour, bitter, umami, sweet and salt) are shown. High and low groups are presented according to the molecular evolutionary rate.

### Archaic origin of LoF variants in taste receptor genes

To further address the origin of taste variants, we explored how modern humans acquired the highly polymorphic taste receptor genes by analyzing the archaic humans Neanderthals (57) and Denisovans (58,59), a likely sister groups to the modern human. Because of the vital function, the same sets of taste receptor genes were present in both Neanderthals and Denisovans, suggesting that they were not subject to any strong multiallelic balancing selection. Virtual genotyping of Denisovan and Neanderthal genomes showed great similarities with current humans, though no sign of pseudogenization (table S4). Intriguingly, even the most frequent mutation rs123321 was not present in Denisovan and Neanderthal genome (table S4). On the basis of these findings we concluded that all of the LoF variants in modern human were acquired after the divergence of the modern and archaic human and that adaptive introgression of population-specific alleles has significantly shaped modern human taste differences (fig. 5).

**Figure 5.**
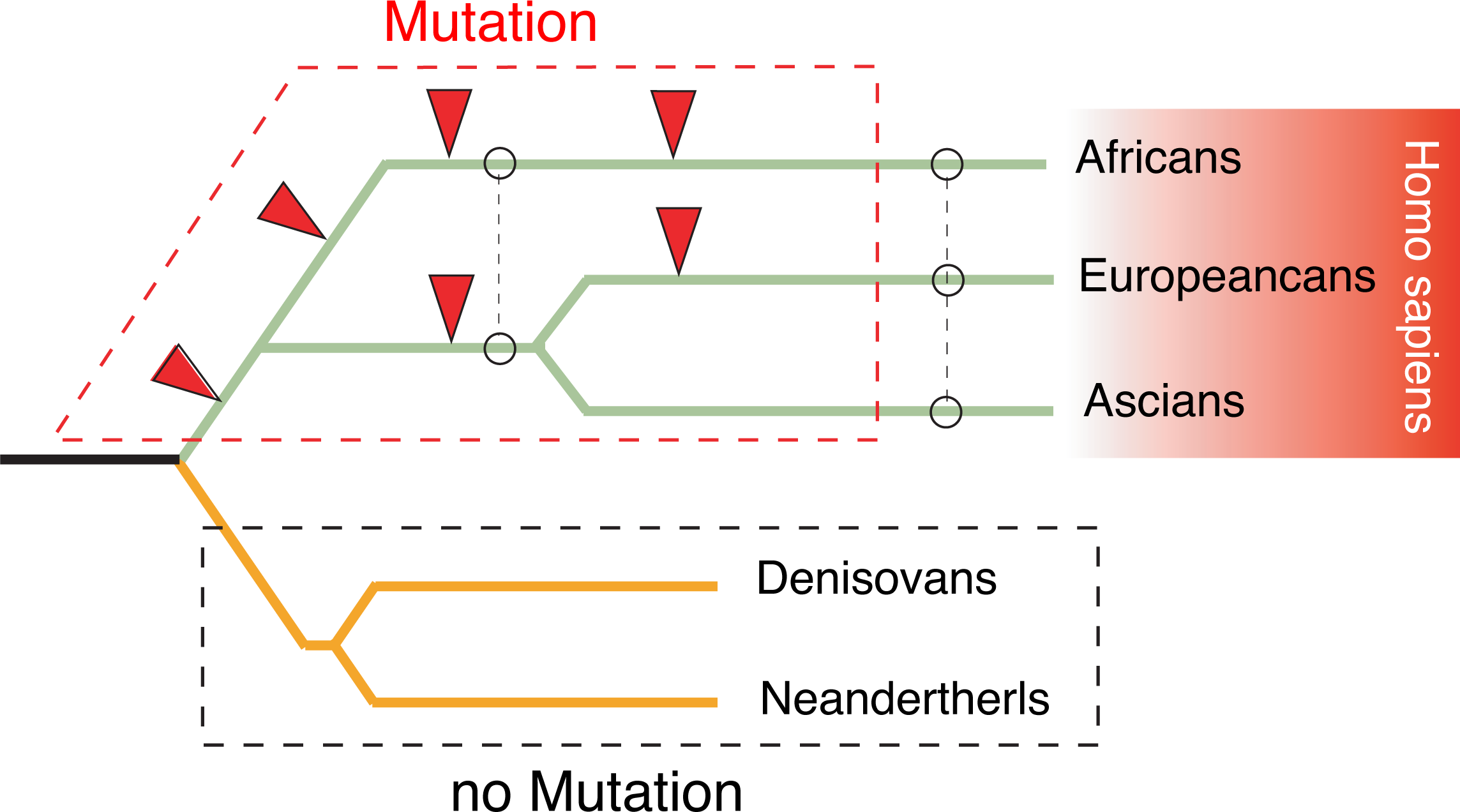
Evolutionary history of loss-of-function variants of taste receptors. Schematic tree shows LoF events of taste receptor genes in modern and archaic humans. Red arrowheads show mutation events. Dashed lines indicate the major introgression between ethnic groups.

### Genetic diagnoses of LoF variants in taste receptors

Finally, we performed preliminary genetic testing for LoF variants in using author’s exome sequences, and examined the effectiveness of the representative list of LoF variants described above. The LoF frequency in taste receptors was extremely high in the human genome, and this case was no exception: three LoF variants, with two in the *PKD1L2* gene and one in the *TAS2R46* gene (Table 1). Although all mutations were heterozygous, we might have a greater risk of vulnerable taste systems with sour and bitter sensations. These mutations followed the pattern of the list shown in Figure 2A and were unexceptional, which supported the effectiveness of this research. Challenges still remain, although our results might yield useful and clinically relevant information for the loss of taste receptors.

**Table 1.** Genetic diagnosis of author’s taste receptors

| Gene name | Transcript ID | Chromosome | Position | dbSNP | Mutation type | Zygoty | ref peptide | Coverage |
| --- | --- | --- | --- | --- | --- | --- | --- | --- |
| *PKD1L2* | NM_052892 | chr16 | 81183325 | rs59980974 | Stop Gain | Hetero | K/* | 28 |
|  |  | chr16 | 81199544 | rs12925771 | Stop Gain | Hetero | R/* | 22 |
| *TAS2R46* | NM_176887 | chr12 | 11214145 | rs2708381 | Stop Gain | Hetero | W/* | 57 |

## Discussion

In 1931, genetic taste variation was first reported as differences in the ability of humans to taste PTC (20). Then many surveys regarding the genetics of taste thresholds were conducted to classify PTC tasters and non-tasters roughly according to their taste acuity by means of dilutions at which the bitter taste was first detected (21,22). It also has long been known that PTC tasters are nonresponsive to the bitterness in *Antidesma bunius* berries, while PTC non-tasters sense bitterness from these berries (60). Although a pile of information about this trait has been accumulated for more than 80 years (20–22), its chromosomal location and causative gene were first reported only in 2003 (13). Over the following ten years, remarkable progress has been made in elucidating the genetic variation of taste ability (29–32), but none of the genetic variants explain the whole spectrum of interindividual differences of taste sensitivity.

In this study we first illustrated the widespread occurrence of a recent rapid decline in the number of functional taste receptors (pseudogenization). Our report is significant because it might raise the hidden possibility that individual taste differences may be driven, in part, by LoF variants in taste receptors. On the other hand we also think that LoF variants in taste receptors not necessarily mean taste impairment because we can consider some scenarios of widespread pseudogenization. Recent study suggested that many hTAS2R receptors share a similar chemical ligands and conversely one agonist can activate several hTAS2Rs, illustrating that hTAS2Rs cooperatively play a common role in toxin avoidance (18,33). Thus some specific taste receptors may have been lost in the human as a result of their functional redundancy. An alternative possibility is that the structural evolution caused by small amino acid substitutions have made the role of chemical receptors less clear, and taste receptors become unnecessary for modern life. Or simply there might be currently no more serious threat to toxins and chemical compounds, or multiple receptors may act as backup for gene inactivation.

Massive loss of taste receptors that we reported is rather unusual, and only a few analogous cases have been described. Another pronounced case is olfaction, which has also undergone a recent rapid decline in the number of functional genes after species differentiation (5,34,35). Olfactory receptors are scattered across the human genome and many genes have already been pseudogenized by nucleotide substitutions (5,34,35). Olfaction and taste receptors cooperatively contribute to flavor, and thus are speculated to have progressed and regressed together during vertebrate evolution (5,34,35). They may have evolved by affecting each other with high expansion rates in addition to selective pressures due to partial functional redundancy (5,34,35). The high-frequency genetic polymorphisms are unique to taste and olfactory receptors in the human genome. This co-evolution could comprise a hitherto unexplored aspect of human genotypic heterogeneity. Changes in these two sensations provide an important clue with regard to future human genetic evolution and may be a major landmark of human evolution.

Recent reports provided some examples that non-synonymous mutations cause interindividual differences of taste sensitivity (32–35). For instance, Africans have higher levels of genetic, geographic and phenotypic diversity at the *TAS2R16* and *TAS2R38* locus (32–35, 61, 62). *TAS2R38* P49A, V296I and A262V have clear effects on PTC bitterness in humans (32–35). In addition non-synonymous polymorphisms in *TAS1R1* and *TAS1R3* also cause variations of sensitivity to L-glutamate (Umami) (29,63). These reports demonstrated that non-synonymous mutations also largely altered bitter and umami taste sensitivity among same populations.

Copy number variants (CNVs) have been reported for *TAS2R43* and the −*45* locus (64). CNVs could result in both overrepresentation and absence of expressed proteins, and have the potential to exert extreme effects on phenotypes. At present, firm conclusions cannot be drawn and more research on genotype–phenotype associations should provide insights into the function of each taste receptor.

My results demonstrated that the LoF variant frequency in taste genes, especially sour- and bitter-related receptors, is likely to be extremely high (fig. 2, 4 and table S2) although not all of the taste receptors were uncovered. Sour and bitter tastes are characteristics of many toxic compounds (65) and have a survival advantage. My result raised the possibility that modern humans lacking sour and/or bitter taste receptors would not seemingly be at a significant disadvantage. In ancient times, meals that may contain toxic substances were life-threatening for our ancestors (66), but because inter-generational or inter-regional exchanges have increased, modern humans may have no need to worry about the risks of daily exposures to toxicants. Consequently, the human intelligence may have driven LoF variants in taste receptor genes. Considering this high mutation rate, “genetically healthy” may be an absolute prerequisite for gastronomists and sommeliers.

I also revealed the country-by-country differences of taste genetic variants. African populations had higher levels of genetic diversity at bitter and fat receptors (fig. 2A and table S3). One possible reason for this might be to increase caloric intake: Africans take traditional low-fat, bitter foods, but must eliminate caloric restriction to alleviate hunger. In this case LoF variants are beneficial in humans. As shown above prevalence of TAS receptors deficiency might be also explained by the hypothesis that modern humans may have no need to worry about toxic compounds after acquiring a wisdom of living.

Recent studies have provided an intriguing example that a loss of sweet receptor genes is widespread among carnivorous species and that taste receptors directly shape feeding behavior (66–68). In contrast to carnivorous mammals, humans might have been losing sour and bitter receptor genes. The recent expansion in number and increased survival rates of modern humans are also speculated to be involved in high-frequency taste LoF variants. Recent research suggested that 90% variants arose within the last 5,000 years (69).

Many testimonies concerning the taste disorders can be collected in immense literatures (70–76). Previously, these disorders could certainly be considered to be related to certain conditions such as sequelae of medications (71), radiation therapy (71,72), smoking (73), chronic systemic illness (74), and nutritional deficiency (75,76). Thus, a strong emphasis has been placed on iatrogenic and secondary taste disorder. In addition, various taste disorders are naturally expected to be curable and reversible (77–79). My results may raise the hidden possibilities of unknown hereditary taste impairment. The mutation frequencies in taste receptor genes could account for the number of different modes (70) of specific taste impairment or human tongue-specific sensory deficits. It is important to realize that a taste disorder, particularly if it is sustained, can have an impact on the quality of a patient’s life (80,8). In general loss of taste can lead to nutritional deficiencies due to anorexia, exacerbations of diabetes, stroke, and hypertension or even contribute to depression (80,81). These symptoms can be particularly serious in older people with chronic illnesses.

Therefore, we hope that my findings might serve as a basis for clinical diagnosis in the near future. Relationships among the taste receptor structures, dietary choices, and the associated metabolic pathways remain unclear. Further biochemical and cell biological studies are currently underway to uncover the total effect of mutations on individual taste sensitivity; they should lead us to a fundamental understanding of the relationship between modern human’s taste and genetic variants.

## Methods

### Analysis of LoF variants from diverse ethnics

We analyzed sequence data from 1000 Genomes projects (http://browser.1000genomes.org/index.html) and NHLBI (http://www.nhlbi.nih.gov/) database. Briefly, these data-sets consisted of high-coverage whole-genome and exome sequence data from diverse ethnic groups, respectively. In addition NCBI dbSNP (http://www.nlm.nih.gov/), UCSC genome browser (http://genome.ucsc.edu/) and HapMap (http://hapmap.ncbi.nlm.nih.gov/) was used as sequencing platform, which were analyzed using an integrated read mapping and variant-calling pipeline to generate our initial catalogue of candidate LoF variants of taste receptor genes. MAF (%) was calculated using computer simulator.

### Multivariate analysis

Hierarchical clustering analysis was performed using the R 3.01 statistical software together with the Rcmdr package. We employed both Ward’s and median algorithms to configure the setting for clustering coefficient. Non-Hierarchical clustering algorithms was based on k-means approach. To separate confused data sets to make distinct classes, principal component analysis (PCA) was also performed on the matrices of spectra from 14+6 ethnic groups. Two-dimensional score plots and loading profiles of the principal components (PC 1 and 2) were applied to visualize the relative contribution of people’s taste preferences to the clustering of the different spectra.

### Evolutionary scenario of LoF variants in taste receptor genes

To estimate how much LoF events in taste receptor genes had occurred before and after the divergence from African origins, we compared alleles among individuals of various ethnic backgrounds Evidence of phylogeny could be based on significant differences in pair-wise comparisons between populations if two groups are significantly different (2-sample test for equality of proportions with continuity correction). The standard hypothesis test is *Η*_0_: π_1_ = π_2_ against the alternative (two-sided) *Η*_1_: π_1_ ≠ π_2_. The pairwise prop test **c**an be used for testing the null that the proportions (probabilities of success) in two groups are the same. It is referred to as a z-test because the statistics looks like

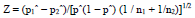

where p^ = (p_1_ + p_2_)/(n_1_ + n_2_), and indices (1,2) refers to the first and second line of the table. In a two-way contingency table where *Η*_0_: p_1_ = p_2_, this should yield comparable results to the ordinary χ 2 test.

### Exome sequencing and bioinformatics analysis

Genomic DNA was extracted from the peripheral blood. Genomic DNA was captured on an Agilent SureSelect Human All Exon V5 71 Mb kit (Agilent) following the manufacturer’s protocols. DNA was sheared by sonication and adaptors were ligated to the resulting fragments. The adaptor-ligated templates were fractionated by agarose gel electrophoresis and fragments of the desired size were excised. Extracted DNA was amplified by ligation-mediated PCR, purified, and hybridized to the capture array using the manufacturer’s buffer. Bound genomic DNA was eluted, purified, and amplified by ligation-mediated PCR. The resulting fragments were purified and subjected to DNA sequencing on the Illumina HiSeq 2000 platform. Captured libraries were sequenced on the Illumina genome analyzer as 100-bp paired-end reads, following the manufacturer’s protocols. Image analysis and base calling was performed by Illumina pipeline versions 1.4 with default parameters. Sequence reads were mapped to the reference genome (hg19) using the Maq program. Identified variants were annotated based on gene name, mutation type and SNP ID. Third-party relatives agreed to a research proposal. The risks associated with participation in this research are thought to be minimum because the frequency of these variants were high in modern populations.

## Data access

Allele information of taste receptors is available from 1000 Genomes projects (http://browser.1000genomes.org/index.html) and NHLBI (http://www.nhlbi.nih.gov/) database.

## Acknowledgments

The author acknowledges Dr. Wolfgang Meyerhof (Nuthetal) for helpful comments on the manuscript. The author would like to thank the 1000 Genomes Project, the NHLBI GO Exome Sequencing Project, and the International HapMap Consortium for the available datasets.

## Contributions

K.F. conceived the idea, carried out data analysis, and wrote the paper.

## References

1. Goldstein, E.B. (2009) Sensation and Perception, 8th ed. Cengage Learning, USA.

2. Chandrashekar, J., Hoon, M.A., Ryba, N.J. and Zuker, C.S. (2006) The receptors and cells for mammalian taste. Nature, 444, 288–294.

3. Yarmolinsky, D.A., Zuker, C.S. and Ryba, N.J. (2009) Common Sense about Taste: From Mammals to Insects. Cell, 139, 234–244.

4. Schiffman, S.S. (1997) Taste and smell losses in normal aging and disease. JAMA, 278, 1357–62.

5. Nei, M., Niimura, Y. and Nozawa, M. (2008) The evolution of animal chemosensory receptor gene repertoires: roles of chance and necessity. Nat. Rev. Genet., 12, 951–963.

6. Drayna, D. (2005) Human taste genetics. Annu. Rev. Genomics Hum. Genet., 6, 217–235.

7. Bachmanov, A.A. and Beauchamp, G.K. (2007) Taste receptor genes. Annu. Rev. Nutr., 27, 389–414.

8. Tordoff, M.G., Shao, H., Alarcón, L.K., Margolskee, R.F., Mosinger, B., Bachmanov, A.A., Reed, D.R. and McCaughey, S. (2008) Involvement of T1R3 in calcium-magnesium taste. Physiol Genomics., 34, 338–48.

9. Tordoff, M.G., Alarcón, L.K., Valmeki, S., Jiang, P. (2012) T1R3: a human calcium taste receptor. Sci. Rep., 2, 496.

10. Laugerette, F., Passilly-Degrace, P., Patris, B., Niot, I., Febbraio, M., Montmayeur, J.P. and Besnard, P. (2005) CD36 involvement in orosensory detection of dietary lipids, spontaneous fat preference, and digestive secretions. J. Clin. Invest. 115, 3177–3184.

11. Sclafani, A., Ackroff, K. and Abumrad, N.A. (2007) CD36 gene deletion reduces fat preference and intake but not post-oral fat conditioning in mice. Am. J. Physiol. Regul. Integr. Comp. Physiol., 293, R1823–32.

12. Montmayeur, J.P., Liberles, S.D., Matsunami, H. and Buck, L.B. (2001) A candidate taste receptor gene near a sweet taste locus. Nat. Neurosci., 4, 492–498.

13. Kim, U.K., Jorgenson, E., Coon, H., Leppert, M., Risch, N. and Drayna, D. (2003) Positional cloning of the human quantitative trait locus underlying taste sensitivity to phenylthiocarbamide. Science, 299, 1221–1225.

14. Nelson, G., Hoon, M.A., Chandrashekar, J., Zhang, Y., Ryba, N.J. and Zuker, CS. (2001) Mammalian sweet taste receptors. Cell, 106, 381–390.

15. Damak, S., Rong, M., Yasumatsu, K., Kokrashvili, Z., Varadarajan, V., Zou, S., Jiang, P., Ninomiya, Y., Margolskee, R.F. (2003) Detection of sweet and umami taste in the absence of taste receptor T1r3. Science, 301,850–853.

16. Bufe, B., Hofmann, T., Krautwurst, D., Raguse, J.D. and Meyerhof, W. (2002) The human TAS2R16 receptor mediates bitter taste in response to beta-glucopyranosides. Nat. Genet., 32, 397–401.

17. Nelson, G., Chandrashekar, J., Hoon, M.A., Feng, L., Zhao, G., Ryba, N.J. and Zuker, C.S. (2002) An amino-acid taste receptor. Nature, 416, 199–202.

18. Meyerhof, W., Batram, C., Kuhn, C., Brockhoff, A., Chudoba, E., Bufe, B., Appendino, G. and Behrens, M. (2010) The molecular receptive ranges of human TAS2R bitter taste receptors. Chem. Senses., 35, 157–70.

19. Huang, A.L., Chen, X., Hoon, M.A., Chandrashekar, J., Guo, W., Trankner, D., Ryba, N.J. and Zuker, C.S. (2006) The cells and logic for mammalian sour taste detection. Nature, 442, 934–938.

20. Snyder, L.H. (1931) Inherited taste deficiency. Science, 74, 151–152.

21. Hartmann, G. (1939) Application of individual taste difference towards phenyl-thio-carbamide in genetic investigations. Ann. Eugen., 9, 123–135.

22. Harris, H. and Kalmus, H. (1949) Chemical specificity in genetical differences of taste sensitivity. Ann. Eugen., 15, 32–45.

23. Mennella, J.A., Pepino, M.Y. and Reed, D.R. (2005) Genetic and environmental determinants of bitter perception and sweet preferences. Pediatrics, 115, e216–222.

24. Hansen, J.L., Reed, D.R., Wright, M.J., Martin, N.G. and Breslin, P.A. (2006) Heritability and genetic covariation of sensitivity to PROP, SOA, quinine HCl, and caffeine. Chem. Senses, 31, 403–413.

25. Knaapila, A., Hwang, L.D., Lysenko, A., Duke, F.F., Fesi, B., Khoshnevisan, A., James, R.S., Wysocki, C.J., Rhyu, M., Tordoff, M.G., et al. (2012) Genetic analysis of chemosensory traits in human twins. Chem. Senses, 37, 869–881.

26. Newcomb, R.D., Xia, M.B. and Reed, D.R. (2012) Heritable differences in chemosensory ability among humans. Flavour, 1.

27. Kiezun, A., Garimella, K., Do, R., Stitziel, N.O., Neale, B.M., McLaren, P.J., Gupta, N., Sklar, P., Sullivan, P.F., Moran, J.L., et al. (2012) Exome sequencing and the genetic basis of complex traits. Nat. Genet., 44, 623–630.

28. Ng, P.C., Levy, S., Huang, J., Stockwell, T.B., Walenz, B.P., Li, K., Axelrod, N., Busam, D.A., Strausberg, R.L. and Venter, J.C. (2008) Genetic variation in an individual human exome. PLoS. Genet., 4, e1000160.

29. Raliou, M., Wiencis, A., Pillias, A.M., Planchais, A., Eloit, C., Boucher, Y., Trotier, D., Montmayeur, J.P. and Faurion, A. (2009) Nonsynonymous single nucleotide polymorphisms in human tas1r1, tas1r3, and mGluR1 and individual taste sensitivity to glutamate. Am. J. Clin. Nutr., 90, 789S–799S.

30. Tepper, B.J. (2008) Nutritional implications of genetic taste variation: the role of PROP sensitivity and other taste phenotypes. Annu. Rev. Nutr., 28, 367–388.

31. Pronin, A.N., Xu, H., Tang, H., Zhang, L., Li, Q. and Li, X (2007) Specific alleles of bitter receptor genes influence human sensitivity to the bitterness of aloin and saccharin. Curr. Biol., 17, 1403–1408.

32. Campbell, M.C., Ranciaro, A., Froment, A., Hirbo, J., Omar, S., Bodo, J.M., Nyambo, .T, Lema, G., Zinshteyn, D., Drayna, D., Breslin, P.A. and Tishkoff, S.A. (2012) Evolution of functionally diverse alleles associated with PTC bitter taste sensitivity in Africa. Mol. Biol. Evol. 29, 1141–1153.

33. Kuhn, C., Bufe, B., Batram, C. and Meyerhof, W. (2010) Oligomerization of TAS2R bitter taste receptors. Chem. Senses., 35, 395–406.

34. Malnic, B., Godfrey, P.A. and Buck, L.B. (2004) The human olfactory receptor gene family. Proc. Natl. Acad. Sci. USA, 101, 2584–2589.

35. Hasin-Brumshtein, Y., Lancet, D. and Olender, T. (2009) Human olfaction: from genomic variation to phenotypic diversity. Trends Genet., 25, 178–184.

36. Bufe, B., Breslin, P.A., Kuhn, C., Reed, D.R., Tharp, C.D., Slack, J.P., Kim, U.K., Drayna, D. and Meyerhof, W. (2005) The molecular basis of individual differences in phenylthiocarbamide and propylthiouracil bitterness perception. Curr. Biol. 15, 322–327.

37. Altshuler, D., Durbin, R.M., Abecasis, G.R., Bentley, D.R., Chakravarti, A., Clark, G. Collins, F.S., De La Vega, F.M., Donnelly, P., Egholm, M., et al. (2010) A map of human genome variation from population-scale sequencing. Nature, 467, 1061–1073.

38. Tennessen, J.A., Bigham, A.W., O’Connor, T.D., Fu, W., Kenny, E.E., Gravel, S., McGee, S., Do, R., Liu, X., Jun, G., et al. (2012) Evolution and Functional Impact of Rare Coding Variation from Deep Sequencing of Human Exomes. Science, 337, 64–69.

39. Menashe, I., Man, O., Lancet, D. and Gilad, Y. (2003) Different noses for different people. Nat. Genet., 34, 143–144.

40. Stahler, F., Riedel, K., Demgensky, S., Neumann, K., Dunkel, A., Taubert, A., Raab, B., Behrens, M., Raguse, J.D., Hofman, T., et al. (2008) A role of the epithelial sodium channel in human salt taste transduction? Chem. Percept., 1, 78–90.

41. Huque, T., Cowart, B.J., Dankulich-Nagrudny, L., Pribitkin, E.A., Bayley, D.L., Spielman, A.I., Feldman, R.S., Mackler, S.A. and Brand, J.G. (2009) Sour ageusia in two individuals implicates ion channels of the ASIC and PKD families in human sour taste perception at the anterior tongue. PLoS One, 4, e7347.

42. Ji, H.L., Zhao, R.Z., Chen, Z.X., Shetty, S., Idell, S. and Matalon, S. (2012) δ ENaC: a novel divergent amiloride-inhibitable sodium channel. Am. J. Physiol. Lung Cell Mol. Physiol., 303, L1013–26.

43. Mummalaneni, S., Qian, J., Phan, T.H., Rhyu, M.R., Heck, G.L., DeSimone, J.A. and Lyall, V. (2014) Effect of ENaC modulators on rat neural responses to NaCl. PLoS One, 9, e98049.

44. Hevezi, P., Moyer, B.D., Lu, M., Gao, N., White, E., Echeverri, F., Kalabat, D., Soto, H., Laita, B., Li, C., et al. (2009) Genome-wide analysis of gene expression in primate taste buds reveals links to diverse processes. PLoS One, 4, e6395.

45. Kusakabe, Y., Shindo, Y., Kim, M.R., Miura, H., Ninomiya, Y. and Hino, A. (2005) cDNA microarray screening for taste-bud-specific genes. Chem. Senses, 30, Suppl 1:i12–3.

46. Fischer, A., Gilad, Y., Man, O. and Pääbo, S. (2005) Evolution of bitter taste receptors in humans and apes. Mol. Biol. Evol. 22, 432–436.

47. Hinds, D.A., Stuve, L.L., Nilsen, G.B., Halperin, E., Eskin, E., Ballinger, D.G., Frazer, K.A. and Cox, D.R. (2005) Whole-genome patterns of common DNA variation in three human populations. Science, 307, 1072–1079.

48. Barreiro, L.B., Laval, G., Quach, H., Patin, E. and Quintana-Murci, L. (2008) Natural selection has driven population differentiation in modern humans. Nat. Genet., 40, 340–345.

49. Lee, W. (2003) Testing the genetic relation between two individuals using a panel of frequency-unknown single nucleotide polymorphisms. Ann. Hum. Genet., 67, 618–619.

50. Garcia-Bailo, B., Toguri, C., Eny, K.M. and El-Sohemy, A. (2009) Genetic variation in taste and its influence on food selection. OMICS, 13, 69–80.

51. Callaway, E. (2012) Evolutionary biology: the lost appetites. Nature, 486, S16–7.

52. Feeney, E., O’Brien, S., Scannell, A., Markey, A. and Gibney, E.R. (2011) Genetic variation in taste perception: does it have a role in healthy eating? Proc Nutr Soc., 70, 135–43.

53. Desor, J.A., Greene, L.S. and Maller, O. (1975) Preferences for sweet and salty in 9 to 15-year-old and adult humans. Science, 190, 686–7.

54. Bacon, A.W., Miles, J.S., Schiffman, S.S. (1994) Effect of race on perception of fat alone and in combination with sugar. Physiol Behav., 55, 603–606.

55. Prescott, J. and Bell, G. (1995) Cross-cultural determinants of food acceptability: Recent research on sensory perceptions and preferences *Trends Food Sci*. Technol., 6, 201–205.

56. Essick, G.K., Chopra, A., Guest, S. and McGlone, F. (2003) Lingual tactile acuity, taste perception, and the density and diameter of fungiform papillae in female subjects. Physiol Behav., 80, 289–302.

57. Green, R.E., Krause, J., Briggs, A.W., Maricic, T., Stenzel, U., Stenzel, U., Kircher, M., Patterson, N., Li, H., Zhai, W., et al. (2010) A draft sequence of the Neandertal genome. Science, 328, 710–722.

58. Reich, D., Green, R.E., Kircher, M., Krause, J., Patterson, N., Durand, E.Y., Viola, B., Briggs, A.W., Stenzel, U., Johnson, P.L., et al. (2010) Genetic history of an archaic hominin group from Denisova Cave in Siberia. Nature, 468, 1053–1060.

59. Meyer, M., Kircher, M., Gansauge, M.T., Li, H., Racimo, F., Mallick, S., Schraiber, J.G., Jay, F., Prüfer, K., de Filippo, C., et al. (2012) A high-coverage genome sequence from an archaic Denisovan individual. Science, 338, 222–226.

60. Henkin, R.I. and Gillis, W.T. (1977) Divergent taste responsiveness to fruit of the tree Antidesma bunius. Nature, 265, 536–537.

61. Bufe, B., Breslin, P.A., Kuhn, C., Reed, D.R., Tharp, C.D., Slack, J.P., Kim, U.K., Drayna, D. and Meyerhof, W. (2005) The molecular basis of individual differences in phenylthiocarbamide and propylthiouracil bitterness perception. Curr. Biol. 15, 322–327.

62. Campbell, M.C., Ranciaro, A., Zinshteyn, D., Rawlings-Goss, R., Hirbo, J., Thompson, S., Woldemeskel, D., Froment, A., Rucker, J.B., Omar, S.A., et al. (2013) Origin and Differential Selection of Allelic Variation at TAS2R16 Associated with Salicin Bitter Taste Sensitivity in Africa. Mol. Biol. Evol., 31, 288–302.

63. Raliou, M., Boucher, Y., Wiencis, A., Bezirard, V., Pernollet, J.C., Trotier, D., Faurion, A. and Montmayeur, J.P. (2009) Tas1R1-Tas1R3 taste receptor variants in human fungiform papillae. Neurosci. Lett. 451, 217–221.

64. Roudnitzky, N., Bufe, B., Thalmann, S., Kuhn, C., Gunn, H.C., Xing, C., Crider, B.P., Behrens, M., Meyerhof, W. and Wooding, S.P. (2011) Genomic, genetic and functional dissection of bitter taste responses to artificial sweeteners. Hum. Mol. Genet., 20, 3437–3449.

65. Meyer-Rochow, V.B. (2009) Food taboos: their origins and purposes. Journal of Ethnobiology and Ethnomedicine, 5, 18.

66. Li, X., Li, W., Wang, H., Cao, J., Maehashi, K., Huang, L., Bachmanov, A.A., Reed, D.R., Legrand-Defretin, V., Beauchamp, G.K. and Brand, J.G. (2005) Pseudogenization of a sweet-receptor gene accounts for cats’ indifference toward sugar. PLoS Genet., 1, 27–35.

67. Zhao, H., Zhou, Y., Pinto, C.M., Charles-Dominique, P., Galindo-González, J., Zhang, S. and Zhang, J. (2010) Evolution of the sweet taste receptor gene Tas1r2 in bats. Mol. Biol. Evol., 27, 2642–2650.

68. Jiang, P., Josue, J., Li, X., Glaser, D., Li, W., Brand, J.G., Margolskee, R.F., Reed, D.R. and Beauchamp, G.K. (2012) Major taste loss in carnivorous mammals. Proc. Natl. Acad. Sci. U S A., 109, 4956–4961.

69. Keinan, A. and Clark, A.G. (2012) Recent explosive human population growth has resulted in an excess of rare genetic variants. Science, 336, 740–743.

70. Fark, T., Hummel, C., Hähner, A., Nin, T. and Hummel, T. (2013) Characteristics of taste disorders. Eur. Arch. Otorhinolaryngol. 270, 1855–1860.

71. Hovan, A.J., Williams, P.M., Stevenson-Moore, P., Wahlin, Y.B., Ohrn, K.E., Elting, L.S., Spijkervet, F.K. and Brennan, M.T. (2010) A systematic review of dysgeusia induced by cancer therapies. Support Care Cancer, 18, 1081–1087.

72. Baharvand, M., ShoalehSaadi, N., Barakian, R. and Moghaddam, E.J. (2013) Taste alteration and impact on quality of life after head and neck radiotherapy. J. Oral. Pathol. Med., 42, 106–112.

73. Krut, L.H., Perrin, M.J. and Bronte-Stewart, B. (1961) Taste Perception in Smokers and Non-smokers. Br. Med. J., 1, 384–387.

74. Hardy, S.L., Brennand, C.P. and Wyse, B.W. (1981) Taste thresholds of individuals with diabetes mellitus and of control subjects. J. Am. Diet. Assoc., 79, 286–289.

75. Heyneman, C.A. (1996) Zinc deficiency and taste disorders. Ann. Pharmacother., 30, 186–187.

76. Field, E.A., Speechley, J.A., Rugman, F.R., Varga, E. and Tyldesley, W.R. (1995) Oral signs and symptoms in patients with undiagnosed vitamin B_12_ deficiency. J. Oral. Pathol. Med., 24, 468–470.

77. Ellul, P., Vella, V. and Vassallo, M. (2007) Reversible dysgeusia attributed to azathioprine. Am. J. Gastroenterol., 102, 689.

78. Kiewe, P., Jovanovic, S., Thiel, E. and Korfel, A. (2004) Reversible ageusia after chemotherapy with pegylated liposomal doxorubicin. Ann. Pharmacother., 38, 1212–1214.

79. Imai, H., Soeda, H., Komine, K., Otsuka, K. and Shibata, H. (2013) Preliminary estimation of the prevalence of chemotherapy-induced dysgeusia in Japanese patients with cancer. BMC Palliat. Care., 12, 38.

80. Sánchez-Lara, K., Sosa-Sánchez, R., Green-Renner, D., Rodríguez, C., Laviano, A., Motola-Kuba, D. and Arrieta, O. (2010) Influence of taste disorders on dietary behaviors in cancer patients under chemotherapy. Nutr. J., 9, 15.

81. Deems, D.A., Doty, R.L., Settle, R.G., Moore-Gillon, V., Shaman, P., Mester, A.F., Kimmelman, C.P., Brightman, V.J. and Snow, J.B. Jr. (1991) Smell and taste disorders, a study of 750 patients from the University of Pennsylvania Smell and Taste Center. Arch. Otolaryngol. Head Neck Surg., 117, 519–528.

